# Information Content of Cortical Recordings During Deep Brain Stimulation in Obsessive-Compulsive Disorder

**DOI:** 10.64898/2026.11.23.768634

**Authors:** Sai P Kate, Victoria L Pipia, Geoffrey W Diehl, Benjamin R Hoenes, Aaron N McInnes, Christi RP Sullivan, Alik S Widge

## Abstract

**Objective:** This study aimed to evaluate network brain activity across both cortical and deep brain (ventral-capsule/ventral-striatum; VC/VS) sites in patients with Obsessive Compulsive Disorder (OCD) that were receiving DBS treatment. We sought to identify relationships between activity in both brain sites, and interactions between them, with evolution of OCD symptomology over treatment. We hypothesized that recordings from cortical sites would carry unique informational content that could predict symptoms and support future adaptive stimulation designs.

**Materials and Methods:** Two patients with severe treatment refractory OCD were implanted with electrodes targeting VC/VS and the supplementary motor area (SMA) bilaterally. Electrodes were connected to Medtronic implantable pulse generators (PC+S or Percept) and could both deliver stimulation and record ongoing local field potential activity. Patients underwent approximately two years of stimulation treatment during which clinical measures of OCD symptoms and brain recordings were regularly collected.

**Results:** Spectral features of VCVS and SMA brain activity had significantly higher entropy (information content) in the signal recorded from the cortical site. A mass univariate partial correlation analysis, controlling for confounders, revealed that cortical and VCVS-cortical interactions were highly represented among the features significantly associated with symptom evolution over the course of treatment (Patient 1: 20 of 30 significant features; Patient 2: 22 of 30 significant features; permutation based false discovery rate correction at q < 0.05).

**Conclusions:** Our findings underscore the importance of capturing cortical activity and VCVS–cortical interactions during DBS research for OCD, with potential implications for the development of more informed and personalized stimulation strategies, including adaptive DBS.

The trial was registered on ClinicalTrials.gov as NCT03184454; https://clinicaltrials.gov/study/NCT03184454.

## Introduction

Obsessive compulsive disorder (OCD) is a psychiatric disorder characterized by recurring, unwanted thoughts (obsessions) and repetitive behaviors or mental acts (compulsions) that an individual feels driven to perform. The behaviors often aim to reduce distress or prevent a feared outcome linked to the obsessions, but can become maladaptive and interfere with daily life. OCD has a lifetime prevalence of 1-3% worldwide ^1,2^ and is associated with significant functional, financial and social impairments and increased mortality ^3–5^.

First-line therapies for OCD include exposure and response prevention (ERP) and pharmacological interventions, such as serotonin reuptake inhibitors (SRIs) ^6^. Unfortunately, some patients do not respond adequately to these treatments, and responders often have residual symptoms ^7^. For this treatment-refractory group, deep brain stimulation (DBS) targeting a striatal/internal capsule site called by multiple names (e.g., the ventral internal capsule/ventral striatum [VC/VS] or anterior limb of the internal capsule [ALIC]) can provide significant benefit. Roughly 60–70% of patients experience a significant reduction in OCD symptoms, validating this target’s therapeutic relevance ^8–10^. However, these outcomes underscore that 30-40% of patients derive minimal benefit from DBS, and even among responders, substantial residual symptoms are common ^9,10^. These limitations highlight a need to refine therapeutic strategies and better understand underlying neurophysiology in order to improve treatment outcomes for individuals with treatment resistant OCD.

OCD is theorized to be a network disorder, meaning that it arises from disrupted communication between different brain regions. The dominant circuit theory of OCD is centered around dysfunction in cortical-striatal-thalamo-cortical (CTSC) circuits ^11–14^. These corticostriatal pathways connect regions of the frontal cortex, basal ganglia, and thalamus, and contribute to reward processing, executive function, and motor and response inhibition ^12,15,16^. Aberrant activity in these so-called “loops” has been reported in OCD, causing increased reinforcement of the repetitive thoughts and behaviors characteristic of OCD ^13,14,17^.

Despite recognition of OCD as a network-based disorder, most prior neurophysiological research has examined individual nodes of the CSTC network in isolation, rather than the interaction between them. Certain DBS systems can record local field potentials (LFPs) from the VC/VS DBS target and connected structures ^16,18–20^. Those recordings may support the discovery of biomarkers for OCD, particularly since they can be collected during both illness and symptom improvement. However, since the pathology of OCD involves both cortical and subcortical regions, it is critical to obtain information from both sets of structures to gain a comprehensive understanding of the neurophysiology of OCD. Cortical LFPs, also known as electrocorticography signals (ECoGs), have unique potential advantages over deep brain LFPs, such as less artefact and greater specificity for sensing ^20^. Additionally, they may provide more precise insight into the cortical activity of the CTSC loops, which in turn allows for a more comprehensive understanding of the pathways of the CTSC loops and their dysfunction in OCD. Therefore, multi-site recording studies may be necessary to capture network-level dynamics that may be crucial in understanding and improving treatment of OCD.

Beyond the potential value of cortical recordings for decoding the neural circuitry of OCD, stimulating cortical regions involved in dysfunctional CTSC loops could have therapeutic benefits for patients with OCD. A particular cortical region of interest for stimulation is the supplementary motor area (SMA), a frontal cortical region implicated in motor planning and habit generation. The SMA is an effective target for transcranial magnetic stimulation (TMS) treatment for OCD^21–23^.

Here we present an exploratory study applying chronic dual-site recording and stimulation in OCD, building on a 2020 report on Patient 1 ^18^. We enrolled two patients with severe, treatment refractory OCD in whom we implanted DBS electrodes bilaterally in the VC/VS and ECoG paddle electrodes bilaterally over the SMA. Using this configuration we collected regular, resting state LFP recordings in both VC/VS and SMA in both patients for over 2 years each. We hypothesize that the cortex harbors unique neurophysiological signatures of OCD that are inherently omitted by a localized VC/VS sensing approach. To evaluate this hypothesis, we conducted two complementary analyses: an assessment of raw LFP signals, and a mass-univariate analysis designed to identify neural features, including power, connectivity, and burst dynamics, that correlate with OCD symptom severity. We then tested whether features derived from VC/VS and cortex/SMA exhibited differential associations with OCD symptom severity.

## Materials and Methods

### Patients

Two patients with severe treatment refractory OCD were enrolled in this study and implanted with electrodes targeting VC/VS and SMA bilaterally. Patients were recruited from specialty neuromodulation clinics. Both gave written informed consent to all study procedures, after verification of capacity to consent. Study procedures were approved by the Institutional Review Boards of Massachusetts General Hospital and the University of Minnesota, and were conducted under the auspices of an Investigational Device Exemption from the US Food & Drug Administration. Importantly, though both of these patients were considered treatment refractory based on their YBOCS scores and clinical evaluations, their individual OCD symptomatology and complexity differed. For specific patient history, see Table 1.

**TABLE 1:** Patient Histories. ECT: Electroconvulsive Therapy. CBT: Cognitive and Behavioral Therapy.

|  | Patient 1 | Patient 2 |
|---|---|---|
| Age at time of Surgery | 20s | 30s |
| Sex | Male | Male |
| Age of OCD Onset | Early Teens | Teens |
| Age of OCD Diagnosis | Unknown | Late Teens |
| Initial YBOCS | 29 | 33 |
| OCD Symptom Presentation | Mental rituals, obsessions with symptoms, counting and symmetries | Somatosensory “not-just-right” feeling, counting, foot-stomping and throat clearing compulsions |
| Other Treatments | VC/VS DBS, CBT, ERP, ECT | TMS, ERP, CBT |
| Previous Medications | Paroxetine, citalopram, mirtazapine, fluoxetine, lorazepam, clomipramine | Escitalopram, fluvoxamine, paroxetine, sertraline, venlafaxine, clomipramine, aripiprazole, lurasidone, olanzapine, quetiapine, risperidone, clonazepam, alprazolam, naltrexone, lamotrigine, valproate, topiramate, dextroamphetamine, lithium, zolpidem, hydroxyzine, trazodone, depakote, buspirone, propranolol |
| Medications at time of surgery | Fluvoxamine, lithium, amphetamine salt, levomefolic acid, lamotrigine, olanzapine, levothyroxine | Gabapentin, lorazepam, olanzapine, imipramine |

### Surgical Implant

Both patients received Medtronic model 3391 deep leads using standard stereotactic surgery with targeting confirmed using MRI and CT co-registration ^24^. VC/VS leads were placed such that the most distal contact (contact 0) was placed within the grey matter of the ventral striatum, 2 mm posterior to the anterior commissure, and the most proximal contact was in the capsular white matter immediately adjacent to the caudate nucleus. We chose the 3391 lead for its greater dorso-ventral span, enabling capture of LFP signals from a broader range of striatal structures. Cortical paddles (Medtronic model 3986/09130, Resume 4-contact paddle lead) were placed on the dorsal surface of the superior frontal gyrus. Targeting was performed visually by the surgeon using cortical landmarks visible during surgery. Final lead placement is illustrated in Figure 1.

**FIGURE 1.**
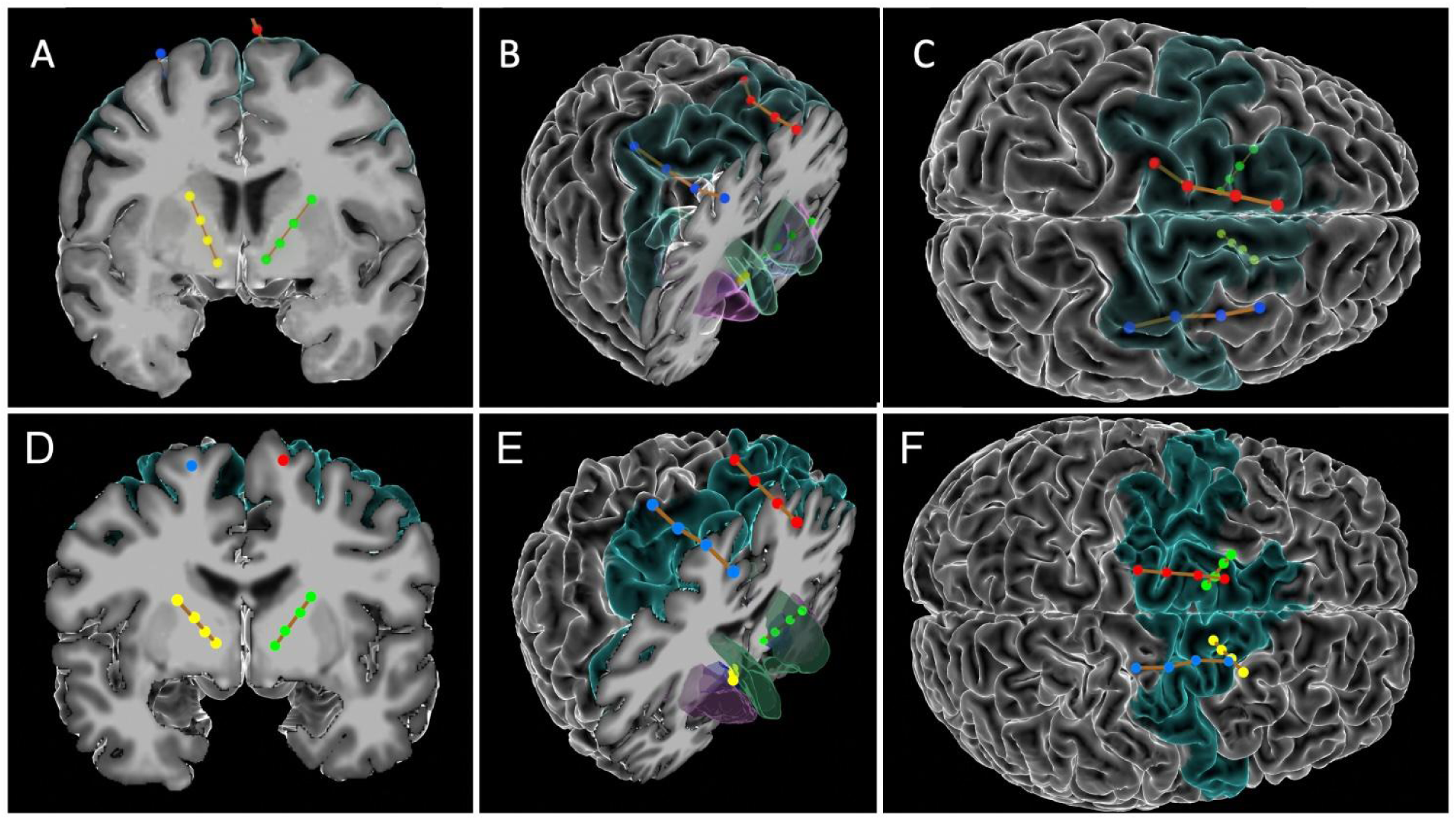
Implanted DBS and paddle contacts for Patient 1 and Patient 2. Images of the implanted DBS and paddle contacts for Patient 1 (top-row; similar visualization from Olsen et al. 2020) and Patient 2 (bottom-row). Left (red) and right (blue) SMA contacts, left (green) and right (yellow) VC/VS are shown with Broadmann Area 6 colored in turquoise. **(A)** Coronal slice showing the position of VC/VS contacts for Patient 1. **(B)** Angled view with cortical coronal slice for Patient 1. Caudate nucleus is colored in green, nucleus accumbens in blue, and putamen in pink for Patient 1. **(C)** Superior view showing the cortical lead positions for Patient 1. **(D)** Coronal slice showing the position of VC/VS contacts for Patient 2. **(E)** Angled view with cortical coronal slice for Patient 2. Caudate nucleus is colored in green, nucleus accumbens in blue, and putamen in pink for Patient 2. **(F)** Superior view showing the cortical lead positions for Patient 2.

In subsequent surgeries Patient 1 received two Medtronic Activa PC+S implantable pulse generators (IPG) and Patient 2 received two Medtronic Percept PC IPGs. Both of these systems are capable of stimulating and recording through both the cortical and VC/VS leads, though precise hardware capabilities differ between the two, as discussed below. Both cortical leads were connected to one IPG and the VC/VS were connected to the other. The IPG was changed because the manufacturer discontinued the PC+S program during the course of the study and the Percept was the only sensing option available to continue.

### Study Design

The intended sample size for this study was n=5, but due to the discontinuation of the cortical paddles, only two patients were implanted ^20^. Both participants progressed through a series of study phases (Figure 2).

**FIGURE 2:**
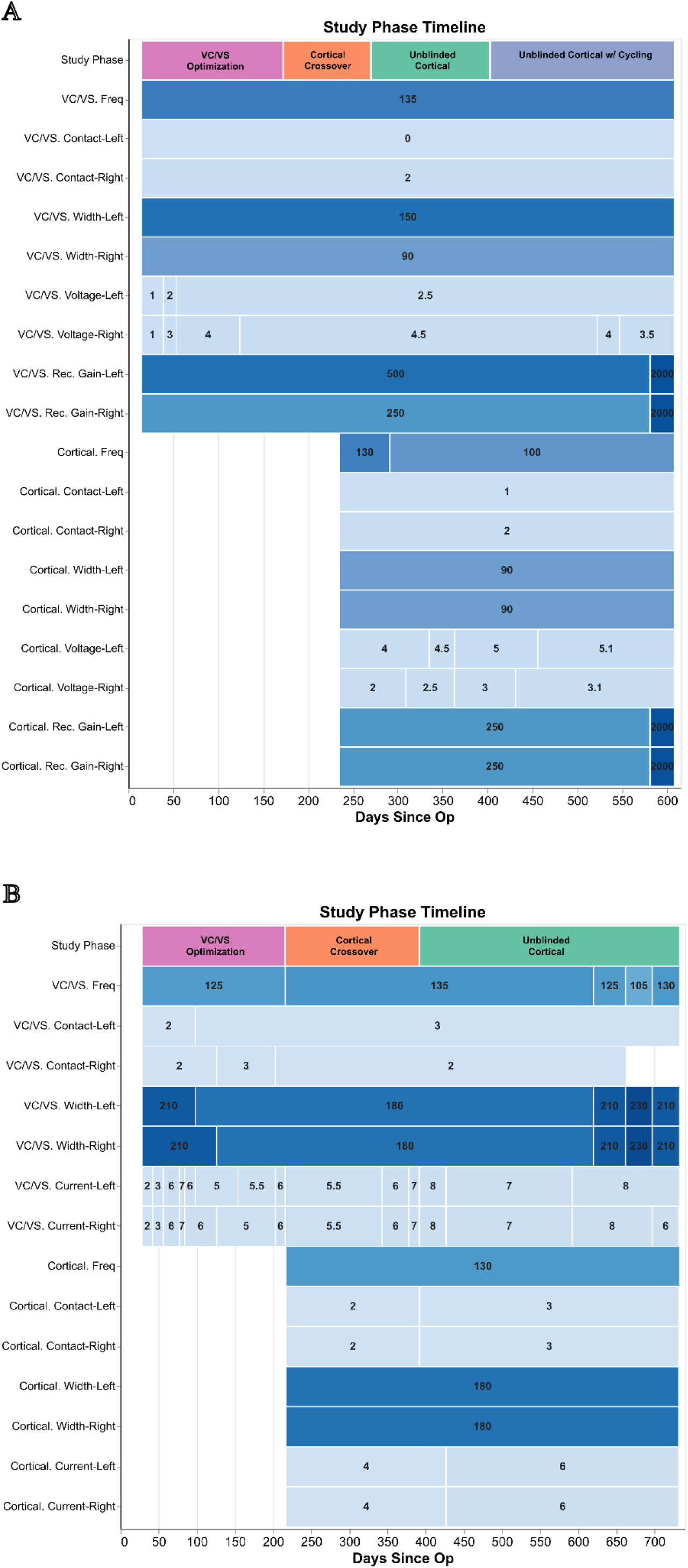
Study timeline and stimulation parameters used throughout the study for Patient 1 (A) and Patient 2 (B). “Width” denotes stimulation pulse width in microseconds. Frequency is measured in Hz. Voltage (Patient 1) is measured in volts and Current (Patient 2) is measured in mA. Some parameters that were configurable in Patient 1 were not available in Patient 2 due to the change in study device.

Study phases were: <u>VC/VS Optimization</u> during which patients received only VC/VS stimulation and stimulation was titrated at regular optimization visits ^24^. In the <u>Cortical Crossover</u> phase cortical leads were then turned on (at a time point unknown to patients or researchers obtaining rating scales). The clinician serving as the DBS programer remained unblinded. When cortical stimulation was turned on by an unblinded programmer, cortical contact and amplitude were optimized and frequency was selected to achieve a slight offset from VC/VS stimulation frequency (e.g VC/VS frequency 135 Hz, Cortical 130 Hz). We theorized that this might effectively disrupt pathological activity in cortico-striatal loops. In the <u>Unblinded Cortical</u> phase the blind was broken on cortical stimulation and patients continued to receive combined Cortical and VC/VS stimulation. Throughout the unblinded phase, patients participated in monthly optimization visits to continue titrating stimulation programming. In Patient 1, although not a formal study phase, there was a period of <u>Unblinded Cortical with Cycling</u> in which stimulation was cycled OFF at night to reduce a subjective sense of activation.

All study components were conducted according to established regulations. The study was approved by the Institutional Review Boards of Massachusetts General Hospital and the University of Minnesota. Both participants provided written informed consent before enrollment. The trial was registered on ClinicalTrials.gov (Identifier: NCT03184454; https://clinicaltrials.gov/study/NCT03184454).

### Stimulation Parameters

Over the two year study period patients underwent three phases of titrating stimulation parameters. In all cases, stimulation parameters were adjusted by a clinician for the purpose of therapeutic benefit. Variables that were adjusted through the study included stimulation contact, stimulation frequency, stimulation pulse width, and stimulation voltage/current. Stimulation in Patient 1 (Activa PC+S) was regulated based on applied voltage (voltage-controlled), while in Patient 2 (Percept PC) it was regulated based on applied current (current-controlled). Note that because of the differences in device, not all parameters were available for both patients. A full description of the stimulation parameters for each patient through the full course of the study is shown in Figure 2.

### Clinical Outcome Measures

Clinical sessions occurred approximately every 2 weeks, where patients received further stimulation titration and clinical outcome data was collected including the Yale-Brown Obsessive Compulsive Scale (YBOCS) ^25^, Montgomery-Asberg Depression Rating Scale (MADRS) ^26^ and Patient Global Impression (PGI) ^27^. Additionally, Patient 2 completed an Obsessive Compulsive Inventory (OCI-R) ^28^ self-report during clinical sessions and remote, self-reported OCD symptom severity scores via the RuneLabs StriveStudy application. These severity scores ranged from 0 (good) to 10 (bad).

To incorporate information from all of the available clinical metrics, and increase the available variability in the clinical data (see Results) we derived a Composite Score. To compute the composite score, each available survey metric (PGI and YBOCS for patient 1; YBOCS, OCI-R and Rune for patient 2) was scaled by its maximum possible score and then summed across all components with a valid scale maximum. The resulting composite score was subsequently z-scored relative to the full data record.

The composite score *C*(*t*) is computed as follows:

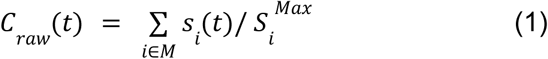

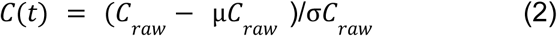

Where *C_raw_*(*t*) is the raw composite score for recording *t*, *M* is the set of survey metrics reported by each patient, and 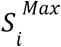 is the maximum possible score for each metric. µ*C_raw_* and σ*C_raw_*mean and standard deviation of *C_raw_* across all recordings, respectively. are

### LFP Recordings

Although the two IPG systems (Activa PC+S vs Percept PC) have differences in their technical capabilities ^29,30^, for the data presented here all recordings were taken using comparable protocols. Recordings were collected approximately once per week, over about 5 minutes of passive rest collected either during clinical visits or at the patient’s home. Stimulation was off during all recordings. Recordings used the “Streaming” feature for PC+S (Patient 1) or the “BrainSense Survey” feature for Percept (Patient 2). Both of these methods collect bipolar recordings, but differ in the contacts measured. In patient 1, recordings were between sites 0 and 2, and in patient 2 they were between 0-2, 1-3, and 0-3. These montages were applied to both the cortical and VC/VS electrodes. For cortical, “0” denotes the most posterior electrode, which is the most distal, because cortical paddles were introduced via the posterior edge of the burr hole. Recordings were collected at either 200Hz (Patient 1) or 250Hz (Patient 2) sampling rate.

Because VC/VS and cortical recording were collected across two independent IPGs, the recording data was not intrinsically time-aligned. At the beginning of each recording session, the clocks of the tablet and both IPGs were synchronized to provide a common temporal reference. The starting time offset between the two recordings was calculated, and the earlier-starting signal was trimmed accordingly so that both recordings began at a common time point.

Previous analyses of data from one patient ^18^ employed a data analysis correction derived from benchtop saline testing of stimulation artifacts produced by the PC+S/Percept systems ^31–33^.

However, comparable testing was not available for the second patient and thus for the current presentation of results these corrections were not used in Patient 1. This change did not make a qualitative difference in the underlying findings from the patient previously described in ^18^.

Following initial data collection, all further analyses were conducted using the MNE-Python package ^34^. The specific neural features of interest are described in the following sections and in Table 2. Because the neural recordings occurred with higher frequency than clinical measures, we downsampled the neural recording metrics by calculating a moving average of neural features from the two weeks preceding each clinical measure.

**TABLE 2:** Overview of features extracted from raw recording data and used for clinical outcome correlation.

| Category | Metric | Brief Description | Patient 1 | Patient 2 |
|---|---|---|---|---|
| Spectral Features | Power | Spectral power containing both components below | Computed for Bilateral Cortical and Striatal Recordings.<br>1 channel per region | Computed for Bilateral Cortical and Striatal Recordings.<br>3 channels per region |
|  | FOOOF’d Power | Spectral power after removing aperiodic (1/f) component |  |  |
|  | Aperiodic Part | The 1/f-like background component extracted by FOOOF |  |  |
| Connectivity Features | WPLI | Measures consistent phase leading/ lagging | Ipsilateral connections only: Cortex: VC/VS<br>1 channel per region = 1 connection | Ipsilateral connections: Cortex and VC/VS<br>Inter-hemispheric connections:<br>R Cortex: L Cortex<br>R VC/VS: L VC/VS<br>3 channels per region = 9 regions |
|  | PLV | Measures consistency of phase differences |  |  |
|  | CIPLV | Corrected imaginary PLV, insensitive to 0-lag connectivity |  |  |
|  | Coherence | Frequency domain connectivity |  |  |
|  | PSI | Directionality |  |  |
| Burst Features | Avg. Cycle Period | Mean period of all cycles | Computed for Bilateral Cortical and Striatal Recordings.<br>1 channel per region | Computed for Bilateral Cortical and Striatal Recordings.<br>3 channels per region |
|  | Avg. Inter-Macroburst Time | Mean time between groups of bursts |  |  |
|  | Number Macro-bursts | Count of detected groups of bursts |  |  |
|  | Number Bursts | Count of individual bursts |  |  |
|  | Avg. Cycle Amplitude | Mean amplitude of all detected bursts |  |  |

### Spectral Analysis

Because all recordings were taken during passive rest, the aligned recording was used as a single analysis epoch, without further subdivision into shorter epochs. Data was decomposed into its time-frequency representation (TFR) using a Morlet wavelet convolution, and then averaged across the time domain to derive the average power at each frequency. TFRs were calculated using 100 wavelet frequencies linearly spaced between 2 and 40 Hz, with the number of cycles increasing linearly from 4 to 7 across these frequencies.This process was repeated for every recording through the duration of the study, in both regions (VC/VS and Cortex) and both hemispheres (right and left). After computing a full TFR and collapsing over time as above, we extracted the mean power in the theta (4-8Hz), alpha (8-12Hz), Alpha/Beta (10-15Hz), low Beta (15-20Hz) and high Beta (20-30 Hz) frequency bands by averaging within those frequencies. Each power spectrum was also parameterized using the FOOOF algorithm ^35^ over 2-40 Hz to separate periodic and aperiodic activity. Periodic band power was then calculated as the mean of the periodic component within the same frequency bands.

We previously proposed that cortical recordings might be more appropriate signals for decoding/predicting clinical symptoms, in part because they may simply be more variable ^20^. To quantify this, we computed the Shannon entropy of the average power across all frequencies, for each recording, hemisphere, and region. Entropy values were pooled across all recordings and study phases and compared between regions within each hemisphere using independent-samples t-tests. Recent reports suggest that entropy, which measures the unpredictability of a signal, may be related to clinical improvement in OCD ^19^. Here, we did not use entropy as a clinical predictor, but as a measure of the potential information content of cortical vs. subcortical recordings. We reasoned that recordings with generally higher entropy were more likely to carry information that could represent clinical biomarkers.

### Burst Activity

Given that brain activity is dynamic, even during passive rest ^36^, we performed a supplemental analysis in which we measured the cycle-by-cycle oscillation activity ^37^. Specifically, we identified the existence of “bursts” of activity (Figure 3) in a particular frequency band (defined by episodes of 5 or more cycles with similar period and amplitude) under the hypothesis that these may represent moments of coherent brain activity which may have relationships to OCD symptomology ^38^. Measuring these bursts also aligns with a recent paper reporting that neural activity becomes less predictable during recovery from OCD ^19^. From each identified burst we extracted relevant characteristics including average cycle period, average amplitude, number of cycles in the burst, and the amount of time between subsequent burst events (inter-burst time). In the event that multiple bursts of the same type were detected with an inter-burst time of less than 100ms, we defined these events as “macro-bursts” as they suggest higher order network structure.

**FIGURE 3:**
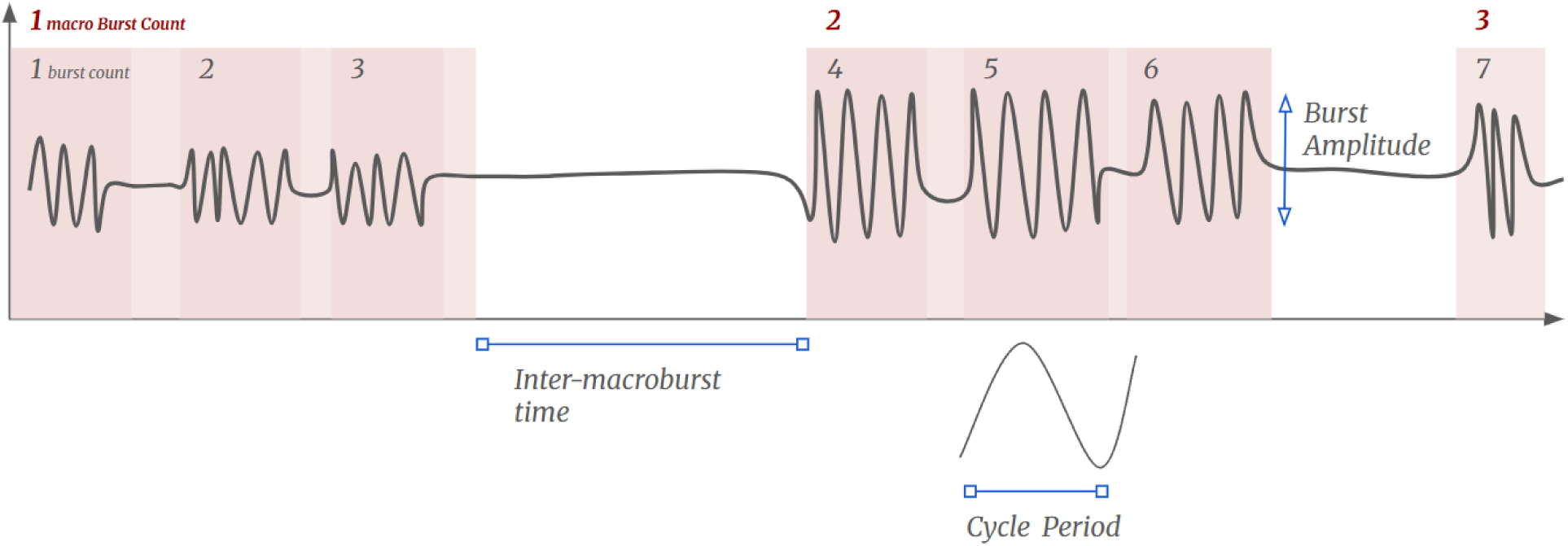
Visualization of burst metric definitions in a time-domain LFP signal. Cycle-by-cycle detection of oscillatory bursts, defined as five or more consecutive cycles with similar period and amplitude, and macro-bursts, defined as clusters of same-type bursts occurring within 100 ms of one another.

### Connectivity Metrics

Time-frequency representations were computed for both brain regions and hemispheres, and for each bi-polar channel pair across relevant frequency bands as outlined in the LFP Recordings and Spectral Analysis sections above. We considered connectivity both between ipsilateral cortical and VC/VS regions, and inter-hemispheric connectivity between like-regions (e.g Left Cortex and Right Cortex). Additionally, for Patient 2, where each lead has three recording channels, each combination of recording channel in region A and region B was included (i.e 9 total channel combinations for each connection investigated). To capture a wide array of activity metrics while remaining agnostic as to any putative best candidate biomarkers we used the python MNE-Connectivity package to calculate weighted-Phase Lag Index (WPLI), Coherence, Phase Locking Value (PLV), Phase Slope Index (PSI), and Corrected Imaginary PLV (CIPLV) for each frequency band.

### Univariate Feature Analysis

To link ongoing neural activity metrics with clinical metrics via our composite score, we performed a mass-univariate analysis. Before conducting univariate analyses, the composite score and neural features were each corrected for potential confounders, specifically days since operation and electrode impedance, by subtracting out the effect of those two variables using ordinary least squares regression. Pearson correlations were then computed between the corrected (residualized) features and composite scores. For each patient, we report the 30 neural features with the strongest correlations to the clinical outcomes.

### Statistical Methods

To assess statistical significance, a non-parametric permutation test was performed in which the null distribution was constructed by applying circular shifts to the corrected composite scores to preserve the temporal structure of the data but disrupt the relationship between specific variations in the features and the clinical outcomes. For each feature, 1,000 permutations were performed and the empirical p-value was calculated as the proportion of permuted correlations with magnitude greater than or equal to that of the observed correlation. In addition to the frequentist p-values, we performed Bayesian Posterior estimation to determine the probability that the effect size exceeded a meaningful threshold (|r| > 0.1). To control the false discovery rate arising from multiple comparisons across high-dimensional features, permutation-derived p-values were adjusted using the Benjamini–Hochberg procedure. Features were considered significant at a corrected threshold of q < 0.05. To compare entropy values between brain regions we used a two sample t-test with significance evaluated at the a < 0.05 level.

We stress that for the neuro-behavioral correlations, we did not perform a cross-validation train/test split, nor did we formally construct a predictive model. The goal of this study was not the development of a predictive approach. Rather, it was a comparison of the informational content and potential predictive power of cortical vs. subcortical originating signals. Correlation on the full dataset satisfies this need.

## Results

### Clinical Outcomes

Clinical outcomes over the course of the study are displayed in Figure 4. Patient 1’s YBOCS score decreased modestly from 27 pre-surgery to <24 by the end of the study. Following the addition of cortical stimulation in week 34 the patient reported significantly improved ability to focus attention away from obsessive thoughts, as evidenced by the decreased PGI. Patient 2 experienced an acute improvement in symptoms as measured by their self-reported Rune score around days 120 - 160 after operation, and a general downward (improvement) trend in YBOCS from 33 pre-surgery to 25 at the end of the study (24.24% reduction). However, Patient 2’s Rune score returned to baseline levels (∼7) after initial improvement and remained elevated for the duration of the study.

**FIGURE 4:**
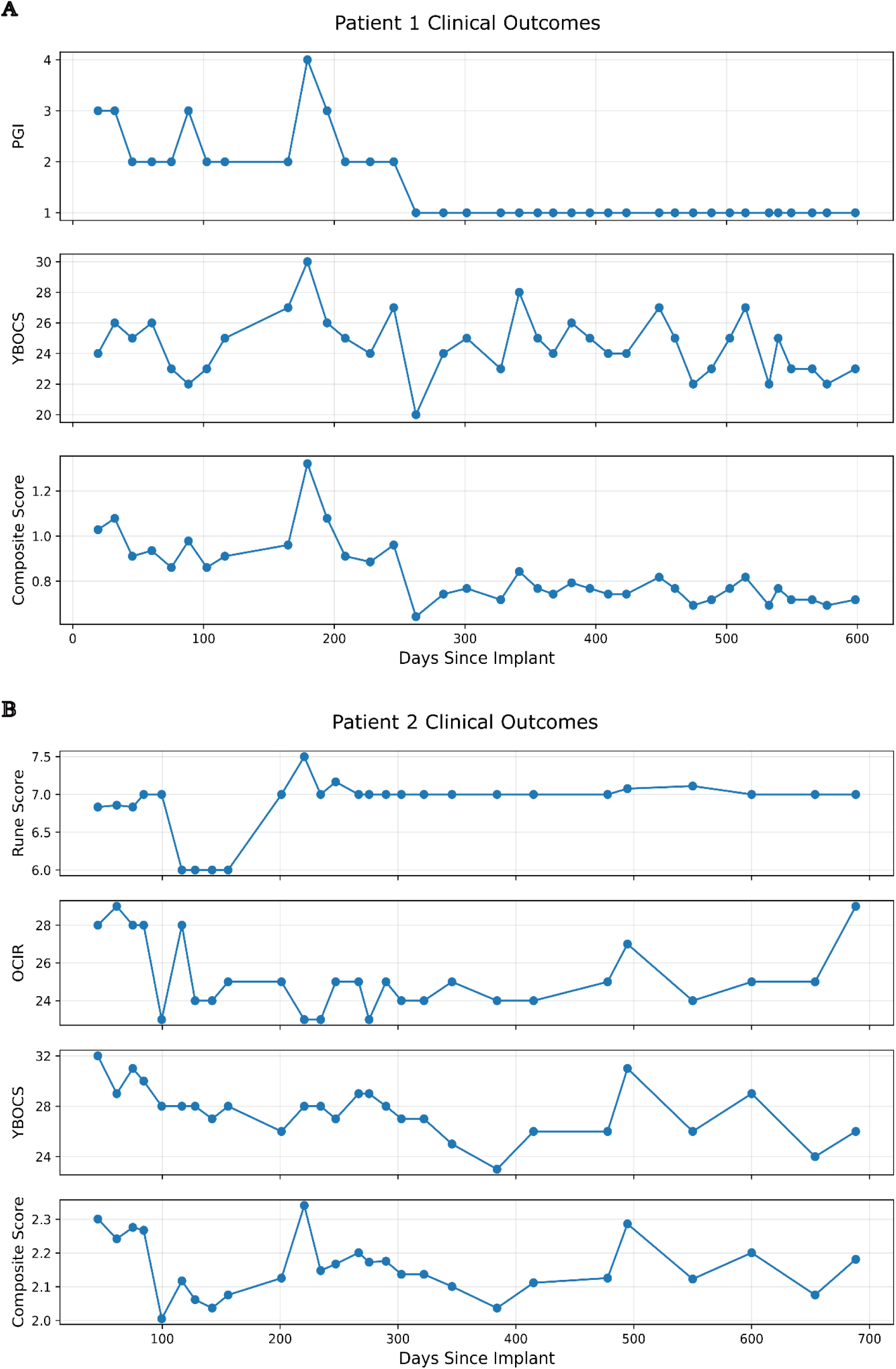
Symptom surveys for Patient 1 and Patient 2 . The X-axis illustrates the duration of the study and the y-axis illustrates the different scores reported by the patient, along with the composite score. Note that the Y-axis range for the two YBOCS plots is different, and that Patient 2 had very limited change on this scale.

For both patients, each individual metric provided a limited picture of overall clinical improvement with some metrics varying non-congruently with others. As such, our composite score attempts to capture any clinical changes within a single unified metric. Furthermore, while both patients exhibited some improvement in their clinical status, neither achieved OCD remission. Thus, while the available data set contains enough variability over time to relate neural activity to clinical metrics, its effectiveness in providing insight into direct OCD related biomarkers is likely limited.

### Cortical and VC/VS LFP Recordings

To evaluate the neural activity from both cortical and VC/VS recordings, we computed the TFR via a Morlet Wavelet convolution and extracted the average spectral power at frequencies ranging from 2Hz to 40Hz. For both patients we observed expected peaks within cortical recordings corresponding to Theta (4-8Hz), Alpha (8-12Hz), and Beta (15-30Hz) oscillations (Figure 5A, B). By contrast, VC/VS recordings were dominated by low frequency activity with a strong peak in the Delta range (4-8Hz). This initial observation suggests that cortical activity contains more overall information across a wider range of relevant oscillatory bands which could be valuable in understanding how neural activity relates to OCD symptomology.

**FIGURE 5:**
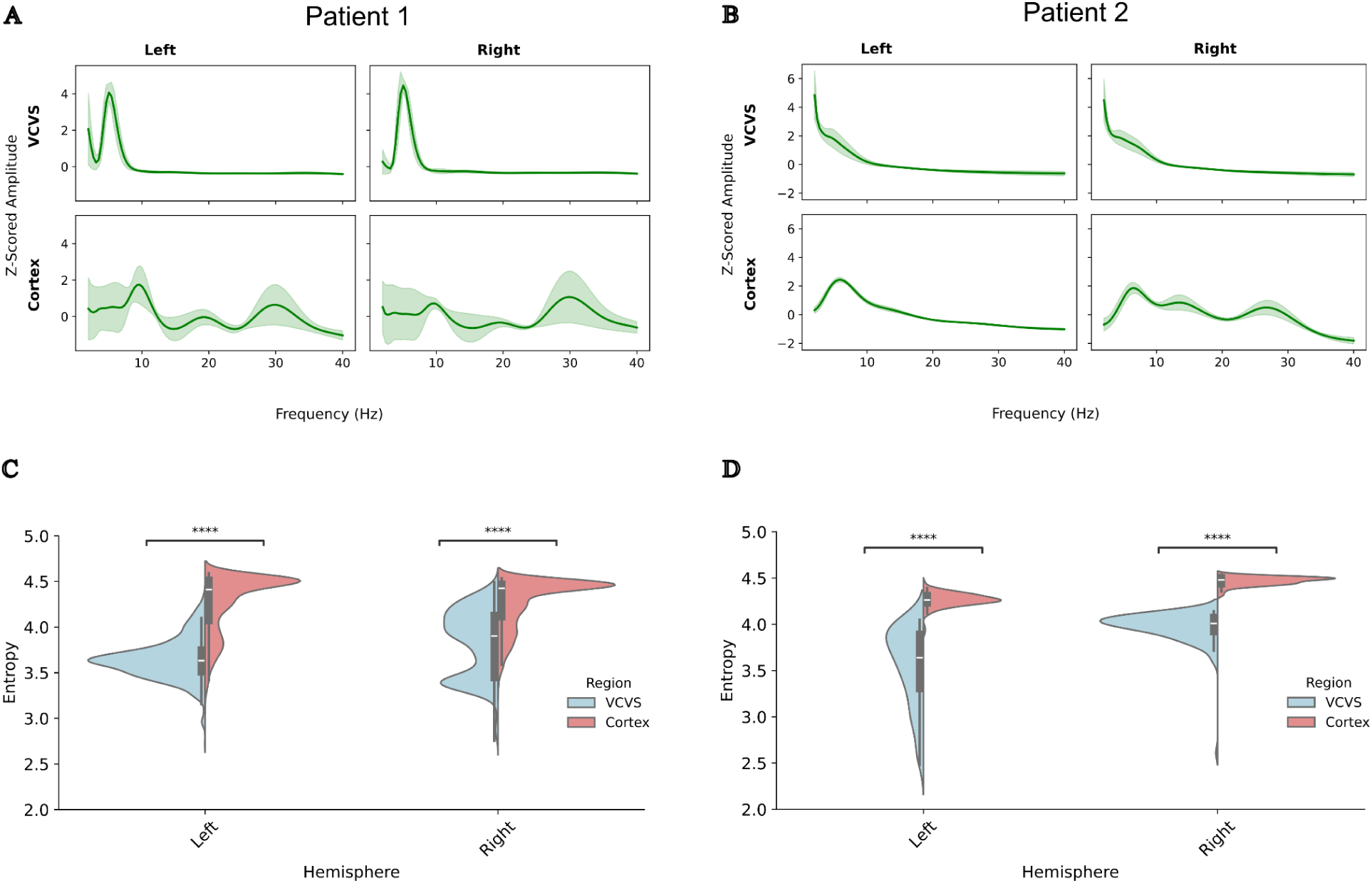
Local field potential (LFP) spectral profiles and entropy across recording regions in two patients. (A, B) LFP recordings in the frequency domain for Patient 1 (A) and Patient 2 (B, across all 3 recording channels), where the solid line represents the z-scored average amplitude at each frequency and the shaded region represents ± one standard deviation across all recordings. (C, D) Entropy was calculated for every recording in Patient 1 (C) and Patient 2 (D), and the distribution of entropies are plotted for VC/VS (light blue) and cortical (red) recordings; t-tests performed between regions (VC/VS vs. cortex) showed significant differences in all cases (**** p < 1e-4).

To directly quantify the level of available information we computed the entropy of the LFP from each recording site in each recording session. For both hemispheres of both patients the entropy of the LFP was significantly higher for cortical recordings as compared to VC/VS recordings (Figure 5). (Patient 1 (Figure 5C): left hemisphere, t(4592) = -77.22, p< 0.01; right hemisphere, t(4592) = -47.57, p< 0.01. Patient 2 (Figure 5D): left hemisphere, t(248) = -18.83, p< 0.01; right hemisphere, t(248) = -31.53, p< 0.01). This elevation in entropy for cortical recordings as compared to VC/VS recordings was present across all study phases (Figure S1) and thus cannot be attributed to any direct effects of electrical stimulation.

Cortical recordings contained distinct spectral peaks across more oscillatory bands (three) as compared to VC/VS recordings (one) and the within-session entropy of the LFP was higher within the cortex. Together these findings suggest that cortical neurophysiology contains more information than a subcortical structure (VC/VS). To determine whether that information was relevant in understanding ongoing OCD symptomology, we next related neural activity to our composite symptom scores.

### Relationship between physiology and clinical scores

To identify VC/VS and cortical neural features associated with symptom severity, we performed a mass univariate analysis. For each region, features were further categorized into intra-hemispheric and inter-hemispheric groups, and grouped by frequency bands within anatomic groups. Features reflecting cross-regional interactions between VC/VS and cortex were designated as VC/VS-Cortex features. After correcting for multiple comparisons, we observed significant associations between symptom score and features in both VC/VS and cortex as well as the VC/VS-Cortex network. Here we present the top 30 features with the strongest correlations with symptom severity for each patient.

For Patient 1 (Figure 6A), burst features in VC/VS and connectivity features in VC/VS-Cortex were the most salient, with VC/VS-Cortex features outnumbering VC/VS features (20 VC/VS-Cortex and 10 VC/VS features of top 30 significant features). In Patient 2 (Figure 6B), cortical features were most strongly represented in the top 30, accounting for 14 of the top 30 features, followed by VC/VS-Cortex and VC/VS features, each represented by 8 features. Among those cortical features, those that measured cross regional communication between the right and left cortex were the most numerous. Across both patients, features involving lower frequency bands such as alpha were more commonly correlated with symptom scores. This is consistent with our spectral analysis where we found elevated power in these same frequency bands. This suggests a preferential importance in evaluating lower frequency bands among patients with OCD, a finding in line with past work ^19,39^.

**FIGURE 6:**
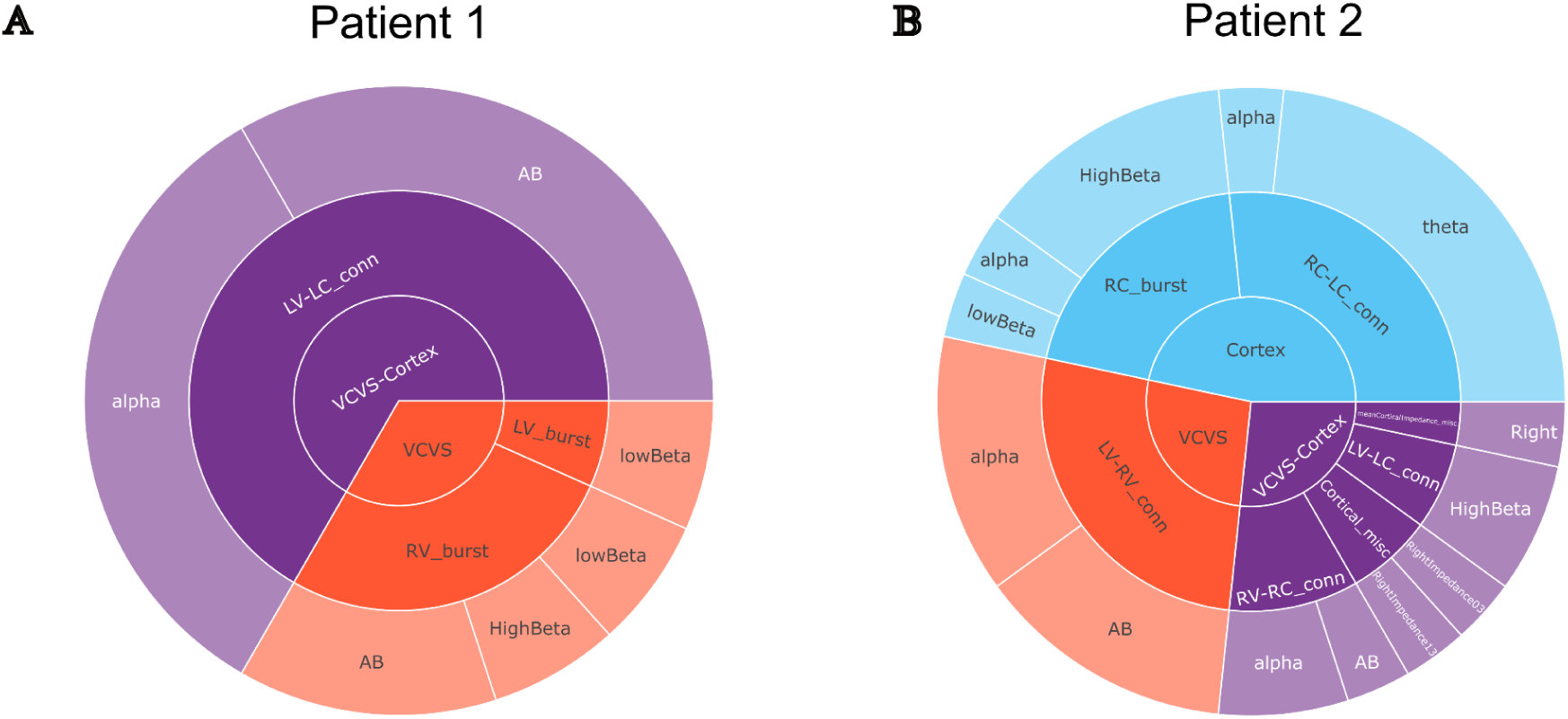
Sunburst plots for Patient 1 and Patient 2 summarizing the top 30 features across both brain regions. Features represent correlations with composite OCD symptom score and are organized according to brain region, feature type, and associated oscillatory frequency band.Segment size reflects the frequency of feature selection LV: Left VC/VS, RV: Right VC/VS, LC: Left Cortex, RC: Right Cortex, Conn: Connectivity based metrics, Burst: Burst based metrics.

## Discussion

Collectively these findings highlight the value of cortical measurements in intracranial studies of OCD, and possibly other psychiatric disorders. In both patients, features requiring cortical recordings carried more information about symptom scores than did features derived only from VC/VS. In Patient 1, inter-regional communication between VC/VS and cortex was prominent while in Patient 2 local and cross-regional activity within cortex itself were significantly correlated with OCD symptom scores. While the precise relationships differed between patients, as might be expected for a heterogeneous condition such as OCD, the observation that cortical activity correlated with symptom scores in both patients suggests a broader utility for measuring cortical activity in this patient population.

Our spectral analysis findings are consistent with prior reports of elevated low-frequency activity at several subcortical stimulation sites, including VC/VS, ALIC, the nucleus accumbens, and pallidum, in patients with OCD, whereas stimulation of the bed nucleus of the stria terminalis (BNST) has been associated with changes in both low- and high-frequency power ^40–44^. In our data, cortical recording sites similarly showed elevated activity across both low- and high-frequency bands. Prior studies have also proposed theta/alpha band as a biomarker of OCD severity ^19,39^. We further found significantly higher entropy in the LFP activity taken from the cortex as compared to VC/VS. We propose that this increased entropy directly points towards a potential value of cortical activity in understanding dynamic OCD symptoms. Cortical activity contains more variation over time, which may be of value in capturing subtle symptom fluctuations. This is in line with recent work highlighting the value of cortical recordings and stimulation in understanding and treating OCD ^45^.

To move beyond local spectral descriptions and probe more complex circuit level interactions relevant to OCD ^46,47^, we implemented a unified analysis framework incorporating power, connectivity, and burst features to characterize regional and hemispheric differences in corticostriatal dynamics. This approach allowed us to assess whether variability between cortical and VC/VS recordings is primarily explained by local oscillatory activity, inter-regional coordination, or transient, non-stationary events. This multiscale perspective is particularly relevant given that corticostriatal circuits likely operate through distributed interactions and temporally structured dynamics that are not fully captured by power alone.

Across this feature space, connectivity and burst-related measures emerged as prominent contributors, with both cortical and VC/VS-derived features showing robust associations with symptom scores. Importantly, the strongest associations were not dominated by a single anatomical source, but were distributed across both regions, indicating that cortical signals are at least as informative as VC/VS signals within this framework. Features capturing VC/VS–cortex interactions were particularly informative, further emphasizing the relevance of circuit-level dynamics in explaining clinical variability.

These findings align with and extend prior work conceptualizing obsessive-compulsive disorder as a disorder of distributed corticostriatal circuitry rather than isolated regional dysfunction within either cortical or striatal nodes ^46^. In particular, theoretical models of OCD emphasize aberrant interactions within cortico-striato-thalamo-cortical loops ^13,48–50^, consistent with our observation that features capturing VC/VS–cortex interactions were strongly associated with symptom severity. Finally, the prominence of connectivity and burst-related features, alongside spectral features, is consistent with recent systems-level approaches that emphasize the importance of transient and interactional dynamics in shaping symptom-relevant brain activity beyond stationary spectral power alone ^19,51^.

More broadly, advances in adaptive DBS have expanded the therapeutic framework beyond the traditional VC/VS target, with increasing emphasis on distributed network activity and the incorporation of cortical signals for biomarker sensing. Our findings provide further support for this direction by demonstrating that cortical recordings are at least as informative as VC/VS recordings, highlighting their potential to enable sense–stimulate closed-loop neuromodulation in which cortical activity guides targeted subcortical stimulation. Given that adaptive DBS in movement disorders has demonstrated that stimulation guided by neural biomarkers, especially cortical signals, can outperform fixed continuous stimulation ^20^, these results provide further rationale for extending similar cortical biomarker-guided neuromodulation strategies to OCD and other psychiatric disorders.

### Limitations

Most notably our study was limited by its small sample size (n=2) and heterogeneity in symptoms and clinical history between the patients intrinsically limiting our ability to identify generalizable biomarkers that could be applied across the broader OCD patient population. The need to change study device similarly added heterogeneity. The subjective nature of OCD symptom metrics and potential lack of alignment across multiple metrics makes robust quantification more difficult. Here, we attempted to reconcile these limitations by deriving a multi-factor symptom score to better capture underlying dynamics of OCD symptoms, though the use of more objective and quantitative metrics will be useful in future work. Finally, we note that local field potential recordings can yield a large number of potential physiologic features which can be related to symptom metrics. This creates a risk of false discovery and over fitting. Thus, we do not claim predictive power, given these substantial limitations. Instead we only aim to highlight the most prominent physiologic features and to make an argument for more extensive use of cortical recordings in future investigations.

## Conclusion

OCD is a network disorder involving cortical and striatal regions. In this exploratory analysis, we used multi-site recordings to demonstrate the feasibility and potential value of capturing these network dynamics in vivo. While the limitations of our data prevent confident, specific biomarker identification, our analysis provides insight into how cortical and striatal signals may jointly inform a more nuanced understanding of OCD pathophysiology. Further work in larger and more diverse patient samples with multi-site recording are needed to characterize OCD pathophysiology and develop therapies targeting the underlying maladaptive network dynamics in OCD and other psychiatric disorders ^52^

## Acknowledgements

We thank Drs. Sarah Olsen and Ishita Basu for their contributions to data collection and clinical care of the patients reported here. We thank the patients themselves for their dedication to the research and their many hours of collaboration.

## Supplementary Figures

The effect shown in the main text, where cortex has higher entropy, is present across all study phases in both patients.

## Financial Support

This study was supported by the NIH (UH3-NS10054801, U01MH116925), the MnDRIVE Brain Conditions Program, and the University of Minnesota Medical Discovery Team on Addictions. Medtronic provided research devices but had no role in the funding, data collection, data analysis, or reporting.

## Author Contributions

VL Pipia, BR Hoenes, AN McInnes, CRP Sullivan, AS Widge designed the study and collected patient data. SP Kate, VL Pipia, GW Diehl, BR Hoenes, AN McInnes, and CRP Sullivan performed data analysis. SP Kate, VL Pipia, and BR Hoenes prepared the initial draft manuscript with intellectual contributions and editing from all other authors. AS Widge secured funding and supervised the study. All authors approved the final version of the manuscript.

## Conflicts of Interests

ASW holds unlicensed patents in the area of deep brain stimulation. He is a founder and holds equity in Resilient Neurotherapeutics.

## Abbreviations

ERP: Exposure and Response Prevention
OCD: Obsessive Compulsive Disorder
DBS: Deep Brain Stimulation
SRIs: Serotonin Reuptake Inhibitors
VC/VS: Ventral Capsule/Ventral Striatum
ALIC: Anterior Limb of the Internal Capsule
CTSC: Cortical-Striatal-Thalamo-Cortical
LFP: Local Field Potentials
ECoGs: Electrocorticography Signals
SMA: Supplementary Motor Area
TMS: Transcranial Magnetic Stimulation
CBT: Cognitive and Behavioral Therapy
ECT: Electroconvulsive Therapy
MRI: Magnetic Resonance Imaging
CT: Computed Tomography
IPG: Implantable Pulse Generators
YBOCS: Yale-Brown Obsessive-Compulsive Scale
MADRS: Montgomery–Åsberg Depression Rating Scale
PGI-I: Patient Global Impression of Improvement
OCI-R: Obsessive-Compulsive Inventory-Revised
TFR: Time-Frequency Representation
WPLI: Weighted-Phase Lag Index
PLV: Phase Locking Value
PSI: Phase Slope Index
CIPLV: Corrected Imaginary PLV

---

**FIGURE S1:**
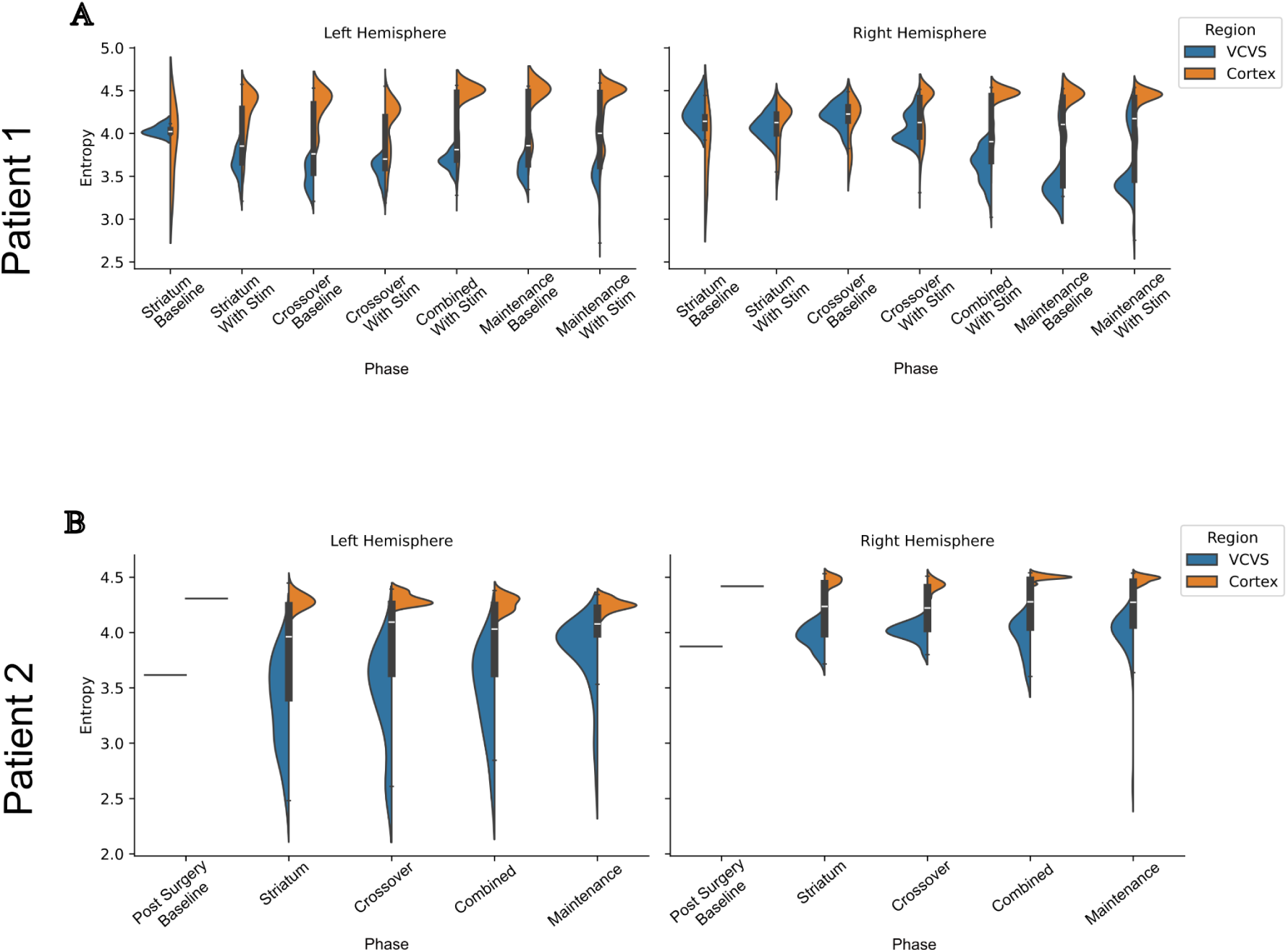
Entropy for each study phase across recordings for Patient 1 and Patient 2. The Entropy of each recording was computed, separated by hemisphere and region (VC/VS or Cortex) and the distribution of recording entropy is shown within a given phase for both Patients. For Patient 2, a single post-surgery baseline was recorded once at the start of the study. This was not available for Patient 1, who had been previously implanted with DBS and had no fully unexposed baseline.

