## Supplementary figures and images for "Information Content of Cortical Recordings During Deep Brain Stimulation in Obsessive-Compulsive Disorder"

### Supplemental Figure 1

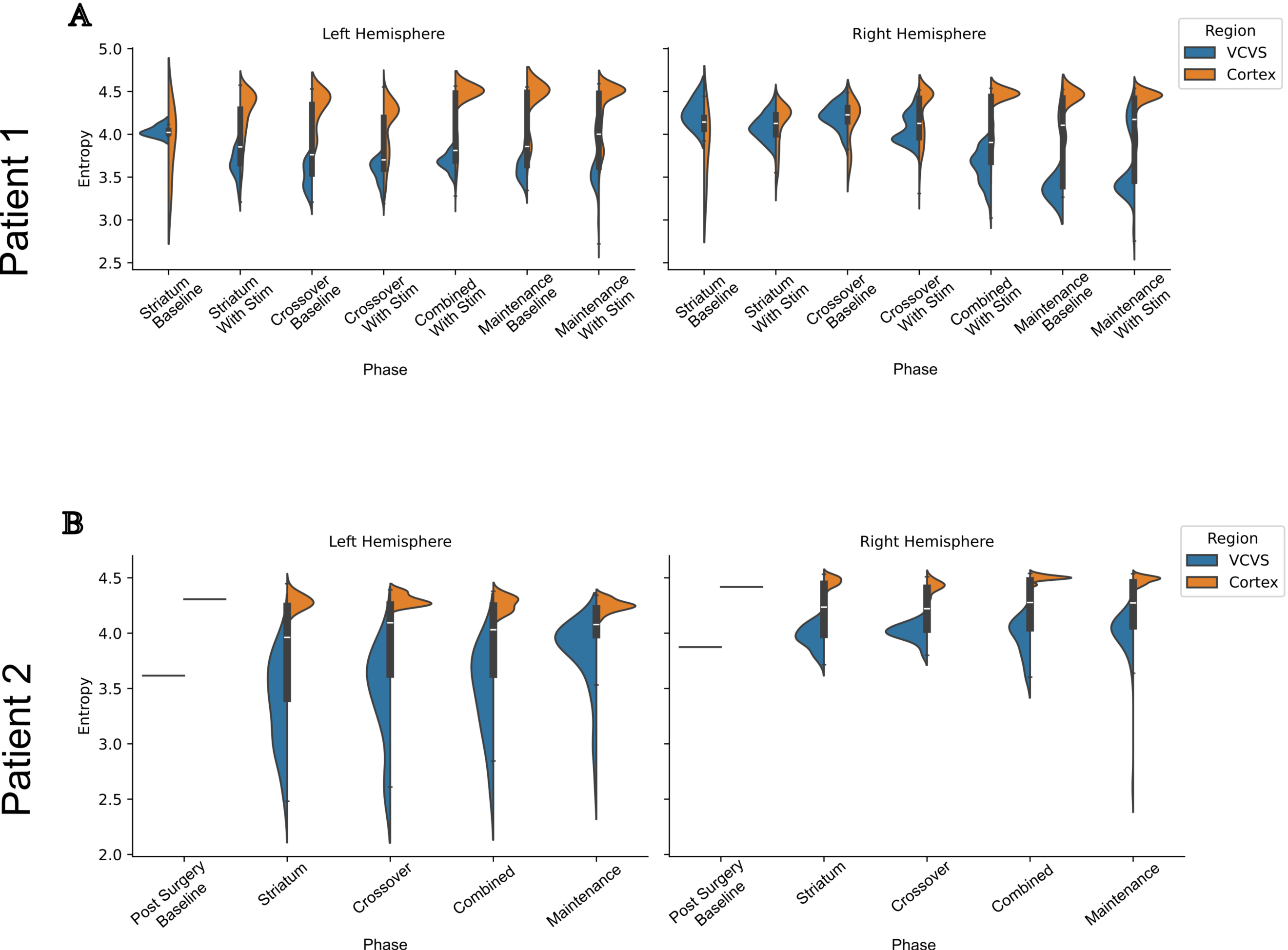
